# EEG-based analyses reveal different temporal processing patterns in aesthetic evaluation

**DOI:** 10.64898/2025.12.12.693903

**Authors:** Limin Hou, Guanghui Zhang, Xiawen Li, Dong Tang, Tiina Parviainen, Fengyu Cong, Tommi Kärkkäinen

**Affiliations:** Institute of Psychological and Brain Sciences, Liaoning Normal University, Dalian, Liaoning, 116029, China; Key Laboratory of Brain and Cognitive Neuroscience, Liaoning Province, Dalian, 116029, China; School of Biomedical Engineering, Faculty of Medicine, Dalian University of Technology, Dalian, 116024, Liaoning Province, China; Faculty of Information Technology, University of Jyväskylä, P.O.Box FI-40014 Jyväskylä, Finland; Center for Mind and Brain, University of California–Davis, Davis, CA, 95618, USA; Department of Physical Education, Shanghai University of Medicine and Health Sciences, Shanghai, 201318, China; School of Foreign Studies, China University of Petroleum (East China), Qingdao, 266580, China; Centre for Interdisciplinary Brain Research, Department of Psychology, University of Jyväskylä, P.O.Box FI-40014 Jyväskylä, Finland; Key Laboratory of Social Computing and Cognitive Intelligence Dalian University of Technology, Ministry of Education, Dalian, Liaoning 116024, China

**Keywords:** Color harmony model, ERP, multivariate pattern analysis, aesthetic evaluation, visual processing

## Abstract

**Background:** Existing studies estimate harmony values using the color harmony model, but how these values are reflected in distinct neural stages (early automatic vs. late evaluative) and shape subjective preferences remains unexamined.

**Methods:** To fill this gap, we recorded behavioral and event-related potential data (N=30 adults, age: 29.20±2.38 years old) to examine whether the predictions of color harmony theory regarding harmonious vs. disharmonious color combinations align with neural activity from a color perception paradigm.

**Results:** Behavioral results showed that two-color combinations received higher aesthetic pleasantness ratings than three-color combinations, while the ratings for color harmony were relatively smaller than for disharmony. Univariate ERP analysis revealed that P1 amplitude for harmonious color was significantly higher than for disharmonious color, particularly when three colors were present. In contrast, the amplitude and latency of P2 were significantly impacted by the number of colors. Additionally, multivariate pattern analysis further demonstrated that neural activity reliably differentiated between harmonious vs. disharmonious color combinations and two-color vs. three-color combinations during both P1 and P2 time window.

**Conclusions:** These findings indicate that harmonious versus disharmonious color combinations subtly influenced early visual processing, while the number of colors used in the design had a stronger influence on aesthetic judgments.

## 1 Introduction

Color harmony refers to the state in which the combination of two or more colors produces a pleasing emotional response and a satisfying visual experience (Burchett, 2002). As a fundamental criterion for assessing the quality and effectiveness of color combinations, color harmony significantly influences user preferences and purchase decisions (Cheng, 2018; Ding et al., 2021). Therefore, many researchers have widely investigated this topic using either behavioral (Chuang & Ou, 2001; Ou et al., 2004a; Schloss & Palmer, 2011) or electrophysiological approaches (Ding et al., 2021, 2023), ranging from self-reported questionnaires and user experience assessments to event-related potential (ERP) studies analyzing neural responses to color stimuli.

To support such investigations, the widely used model is Moon and Spencer’s classical aesthetics measurement theory that offers a widely applied quantitative framework for evaluating color harmony (Moon & Spencer, 1944a, 1944b). Based on this theory, the aesthetic values of multi-color combinations are computed and ranked to identify those with higher harmony scores. The previous studies found that harmonious combinations are consistently rated higher than disharmonious ones by observers, with their subjective evaluations closely aligning with the aesthetic values predicted by the measurement model (Hsiao & Yang, 2017; Shih-Wen et al., 2017). Regarding the influence of the number of colors in combinations, although individuals generally prefer harmonious multicolor schemes (Palmer & Schloss, 2010), some studies observed that increasing the number of colors (e.g., from two to three) leads to a decline in perceived harmony and a rise in visual cognitive load (Ou & Luo, 2006).

In the field of visual neuroscience, color is probably the dimension that has received least attention, but there are still established line of research focusing on the neural mechanisms underlying color perception. Research has demonstrated that different color combinations substantially influence the latency and amplitude of ERPs, particularly at early perceptual and later cognitive stages. For example, the P1 component, which reflects the initial processing of visual information, is primarily influenced by physical attributes such as color or contrast (Hillyard & Anllo-Vento, 1998). Similarly, the N1 component is closely linked to selective attention and feature recognition, and its amplitude variation is thought to reflect later stages of selective processing (Eimer, 2014; Luck et al., 1990). At more advanced stages of cognitive processing, the P2 component and particularly the posterior P2 has been found to be strongly associated with the perceptual aspects of visual evaluation (Herbert et al., 2006; Scott et al., 2009). This component is considered to represent higher-order perceptual processing modulated by attentional mechanisms (B. Liu et al., 2012; Luck & Hillyard, 1994; Omoto et al., 2010).

Although previous studies have investigated the aesthetic effects of harmonious color schemes and provided important theoretical and behavioral insights into color harmony perception (Ou & Luo, 2006), the relationship between theoretically derived harmony values and corresponding neurophysiological responses has yet to be established. In addition, existing research has primarily focused on harmonious combinations, while the cognitive and neural mechanisms underlying the differential processing of harmonious vs. disharmonious color schemes remain largely underexplored (Ding et al., 2022; K.-R. Li et al., 2022). Furthermore, existing studies on color harmony theory tend to emphasize emotional and evaluative aspects, rather than differentiating between early automatic perception and later cognitive evaluation (Hsiao et al., 2008; Hsiao & Yang, 2017) or examining how the number of colors modulates processing across distinct cognitive stages (Ding et al., 2023).

The aim of this study is to investigate whether the harmony and disharmony values of multicolor combinations, as computed based on color harmony theories, are associated with distinct neural mechanisms across different stages of cognitive processing. Unlike previous studies that primarily focused either on emotional impressions or color quantity, this study considers both harmony values and color numbers. It examines how these two factors jointly influence visual and aesthetic processing. Specifically, we measured the amplitude and latency of P1, P2, and N1 ERP components to examine whether the theoretically predicted harmonious and disharmonious two-color and three-color combinations involve both early visual processing and aesthetic evaluation. To further evaluate the significance of activation in these three time-windows for perceptual evaluation of color harmony, we employed multivariate pattern analysis (MVPA) to classify ERP data at each time point across entire electrodes. By integrating univariate and multivariate analysis, this study seeks to provide deeper insights into how color harmony and different number of colors shape neural responses, thereby enriching our understanding of how the brain processes aesthetic and perceptual attributes.

This study hypothesizes that color harmony and the number of colors selectively influence different stages of neural processing—specifically, that color harmony primarily affects early components (P1/N1), while the number of colors modulates later components (P2).

## 2 Materials and methods

### 2.1 Participants

A total of Thirty-five healthy, Chinese undergraduate and graduate students from the University of Jyväskylä, Finland, with normal or corrected-to-normal vision participated in the study. None of the participants reported any history of neurological or psychiatric disorders. ERP data from five participants were excluded because of excessive EEG artifacts with fewer than 70% of trials remaining after rejection, leaving a final sample of 30 participants (29.20 ±2.38 years old; range = 24-34 years old), including 14 males and 16 females.

### 2.2 Experimental design

#### 2.2.1 Experimental sample preparation

Based on the painting and panel processing technologies for automotive bodies, the vehicle body is divided into six major regions. By randomly selecting two or three regions, two-color and three-color schemes for automotive body layouts can be generated. According to an analysis of existing market product painting schemes and concept car body color layouts, the two-color combinations yielded seven design layouts, while the three-color combinations resulted in nine design layouts. To identify the most user preferred two-color and three-color automotive body color layouts, the seven two-color and nine three-color layouts were rendered using neutral colors to minimize the influence of specific colors on user preferences (Details of the layout scheme design and user preference survey, including the selection process and participant demographics, are provided in Supplementary Material). A preference survey was conducted via questionnaires, collecting a total of 118 valid responses. The survey results were analyzed to determine the most popular two-color and three-color automotive body color layout designs, as shown in Fig.1a.

The Munsell color system was adopted to achieve a systematic representation of colors, and the “tone-map” concept from the PCCS color system was applied to construct a comprehensive foundational color database. The Munsell color system characterizes every perceivable color using three dimensions: hue, value, and chroma. It identifies 10 fundamental hues—R (red), YR (yellow-red or orange), Y (yellow), GY (green-yellow), G (green), BG (blue-green), B (blue), PB (purple-blue), P (purple), and RP (red-purple). Each of these hues is further subdivided into four increments (2.5, 5, 7.5, and 10), designated by a numerical prefix, resulting in a total of 40 distinct hues. White, black, and grey are not considered hues in the Munsell system.

For this study, 33 colors were selected to form the color palette. (Fig. 1b). First, 10 fundamental hues at Hue=5 were selected from the Munsell color system. Each hue was then developed into three variants (deep, vivid, and bright) using the PCCS color system, resulting in 30 chromatic colors. Further, three commonly used automotive colors: black, white, and gray were incorporated to provide a comprehensive representation of colors for analysis.

**Fig. 1.**
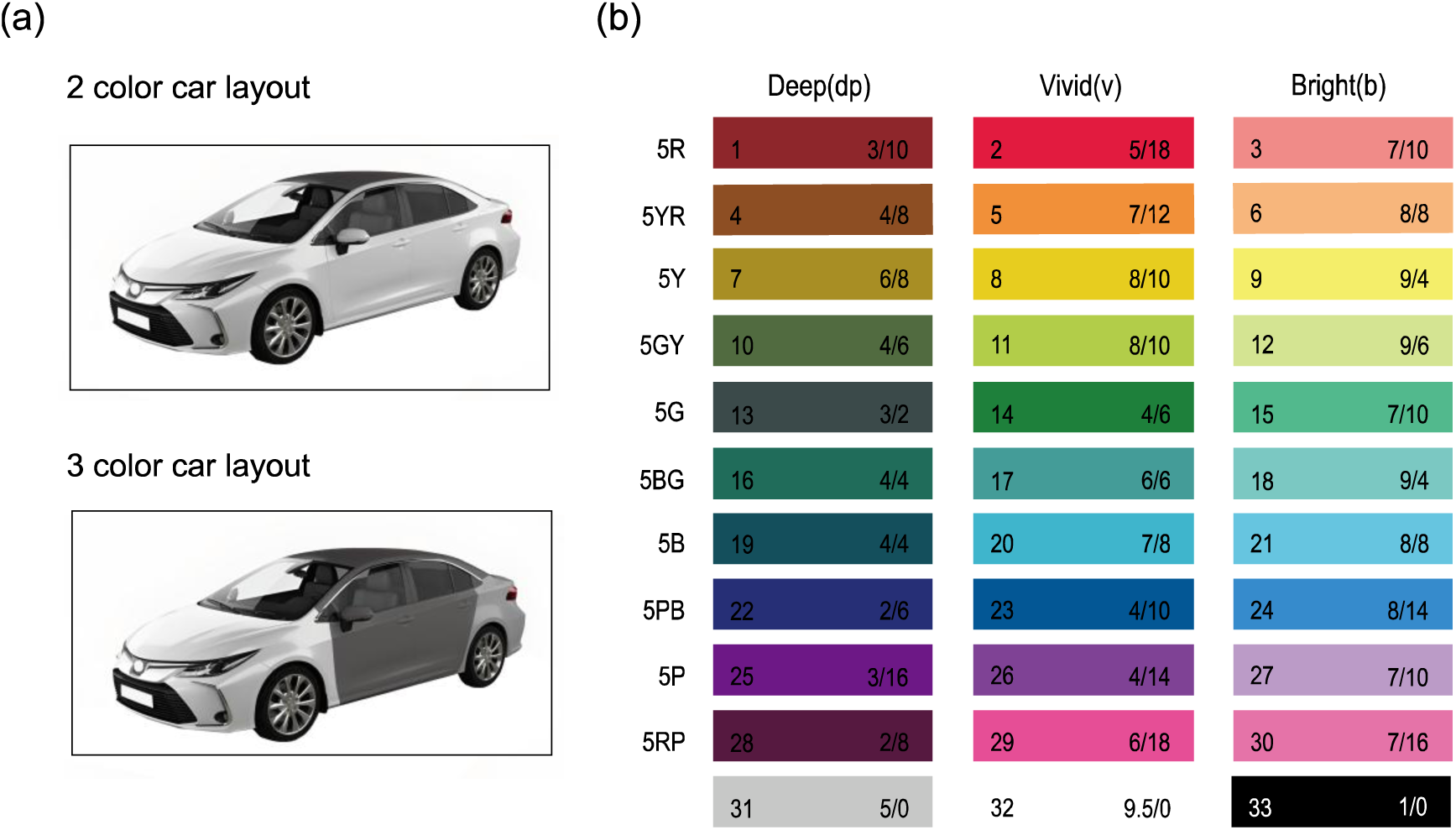
(a) Schematic layouts of two-tone and three-tone car designs. (b) The color palette comprises 33 colors used in this study. R is red, YR is yellow-red or orange, Y is yellow, GY is green-yellow, G is green, BG is blue-green, B is blue, PB is purple-blue, P is purple, and RP is red-purple.

**Fig. 2.**
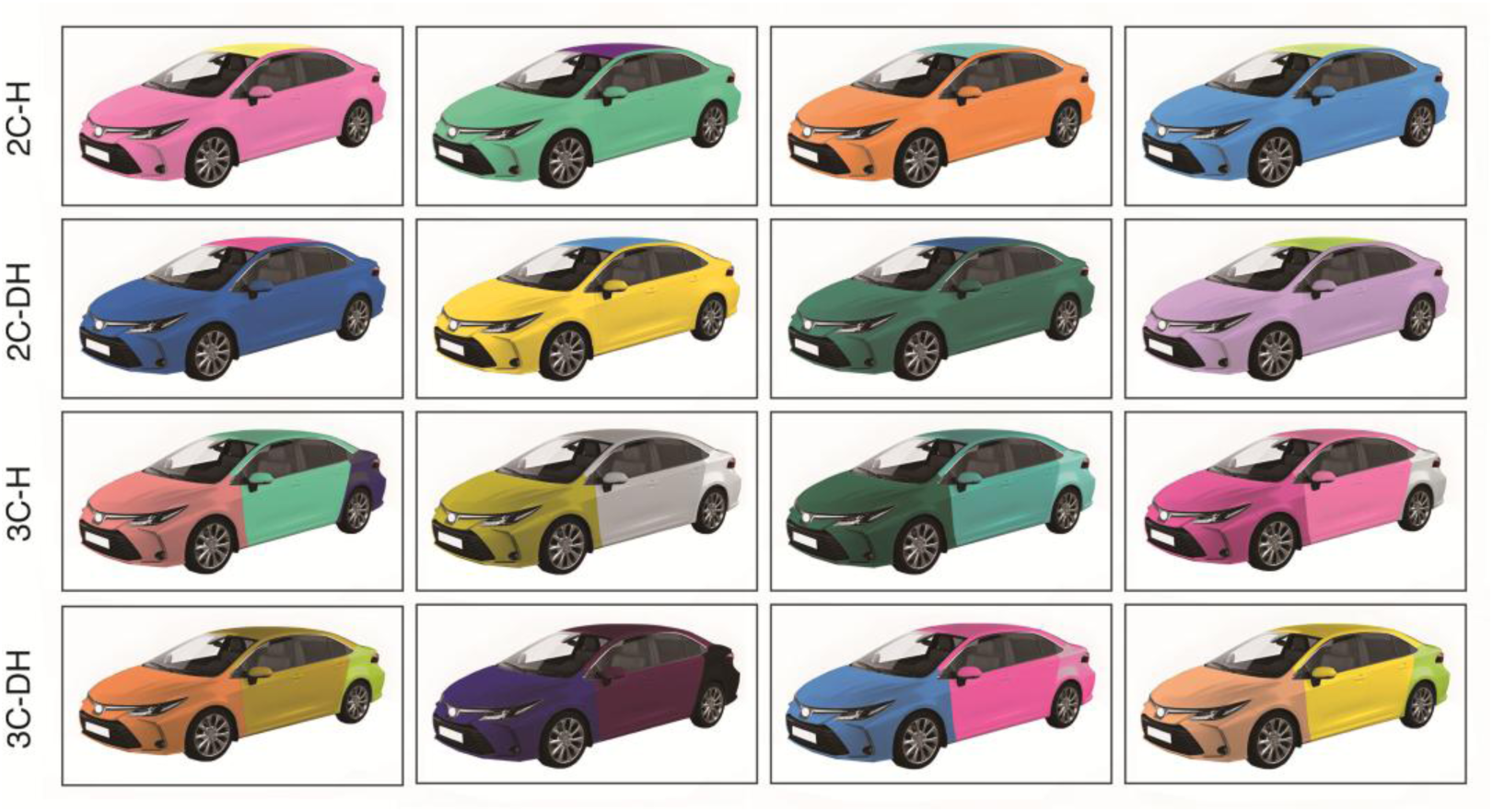
Illustrations of the four experimental stimuli (2C-H, 2C-DH, 3C-H, 3C-DH). Thus, the experiment included two independent factors: (a) scheme complexity (two levels: two-color vs. three-color) and (b) color harmony (two levels: harmony vs. disharmony), resulting in four experimental conditions: 2C-H (two-color harmony), 2C-DH (two-color disharmony), 3C-H (three-color harmony), and 3C-DH (three-color disharmony) schemes. Each condition comprised 100 combinations, yielding a total of 400 image stimuli. Fig. 2 illustrates examples of both harmonious and disharmonious two-color and three-color vehicle designs.

#### 2.2.2 Stimuli

The experimental stimuli were created using Adobe Photoshop (version 24.7.0) and comprised a total of 33 colors. Two-color and three-color combinations were randomly selected and applied to vehicle body designs. Based on the combination formula, 528 possible two-color combinations and 5456 possible three-color combinations were calculated. The color harmony values of these combinations were ranked from highest to lowest based on the color harmony rule proposed by Moon & Spencer (Moon & Spencer, 1944b, 1944a). Since many colors shared the same aesthetic value, combinations with differing aesthetic values were avoided as much as possible when selecting stimuli. To examine the consistency between color harmony theory and EEG data, the 100 combinations with the highest aesthetic values (most harmonious) and the 100 combinations with the lowest aesthetic values (least harmonious) were selected as experimental stimuli for both two-color and three-color schemes.

### 2.3 EEG recording and preprocessing

EEG data was recorded using a 128-channel net (HydroCel Geodesic Sensor), a high-impedance Net Amps 400 amplifier (Electrical Geodesics Inc., Eugene, OR, USA), and Net Station 4.5.7 software (Electrical Geodesic Inc., Eugene, OR, USA) at a sampling rate of 1,000 Hz. The online filters were set up from 0.1 to 250 Hz, and the online reference electrode was placed at a vertex electrode (Cz). The impedance for each electrode for each subject was below 50 kΩ.

Participants were tested individually in a quiet, dimly lit room. The experimental stimuli were presented on a white background using E-Prime 2.0 software (Psychology Software Tools, Inc., Sharpsburg, PA, USA). The experiment consisted of four condition blocks (2-color vs. 3-color × harmony vs. disharmony), presented in a randomized order. Each block comprised 100 fully randomized trials. Prior to the formal experiment, participants completed a practice session of 20 trials (5 images from each condition) to familiarize with the procedure. Each trial began with a black fixation cross (‘+’) displayed for 1000 ms, followed by the presentation of a stimulus image (a 2C-H/2C-DH/3C-H/3C-DH car) for 1000 ms at the center of the screen. After the stimulus presentation, a blank screen was shown for 500 ms, after which a 7-point Likert scale appeared at the center of the screen. The scale remained present until participants provided their response. Participants were instructed to rate the valence of each stimulus (two-color or three-color car) on the 7-point Likert scale, where 1 corresponded to ‘very unpleasant’ and 7 to ‘very pleasant’. The experimental procedure and stimulus paradigm are illustrated in Fig. 3.

**Fig. 3.**
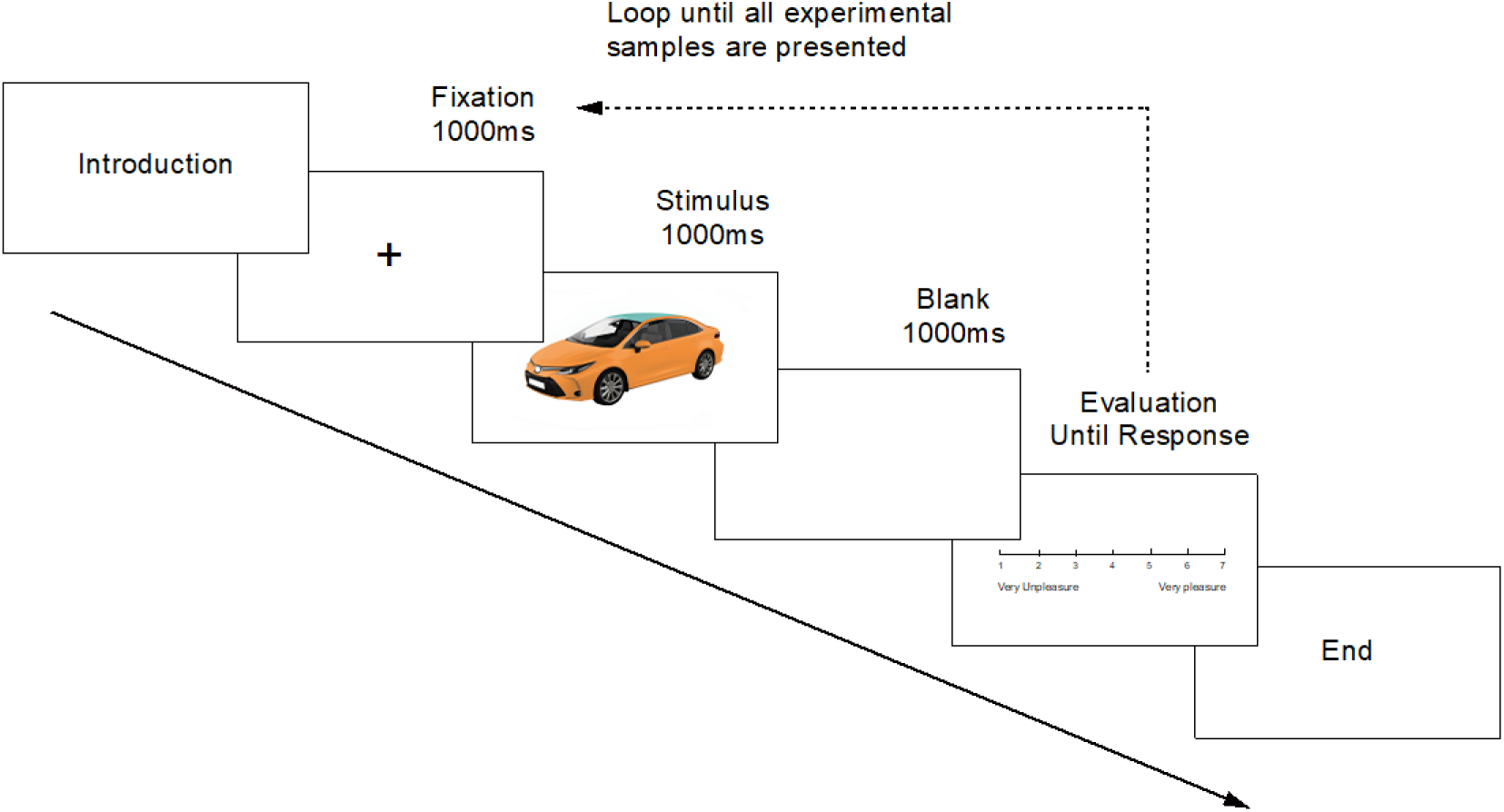
An example of a single trial in the experiment. Each trial began with a fixation cross displayed for 1000 ms, followed by a 1000 ms presentation of the stimulus. After a 500 ms blank screen, participants were prompted to rate the stimulus on a 7-point Likert scale, indicating their level of pleasantness (1 = very unpleasant, 7 = very pleasant).

The following preprocessing steps were included for the offline analyses, which were conducted using EEGLAB 2024 (Delorme & Makeig, 2004). First, the EEG data were filtered using a notch filter to remove 50 Hz line noise. A high-pass FIR filter at 0.1 Hz and a low-pass FIR filter at 30 Hz, with default parameters in EEGLAB, were applied to the EEG data to remove high- and low-frequency noise (Zhang, Garrett, & Luck, 2024a, 2024b). Second, the filtered data were resampled to 500 Hz and re-referenced to the average reference, excluding the EOG channels (8^th^, 25^th^, 126^th^, and 127^th^ electrode channels) (Jia & Tyler, 2019). Next, independent component analysis (ICA) was employed using the ICASSO software to eliminate artifacts by removing independent components (ICs) related to eye blinks and movement activities (Zhang, Garrett, Simmons, et al., 2024). Bad channels with significant noise were interpolated using the spherical spline method. Afterwards, the EEG data were segmented from 200 ms before stimulus onset to 800 ms after stimulus onset. Baseline correction was applied to each epoch by subtracting the mean voltage of the pre-stimulus period (−200 ms to 0 ms) from each time point. The trials were discarded if the amplitude exceeded ±80 μV, resulting in 91.85% ± 6.01% of trials being retained per participant.

### 2.4 Univariate ERP analysis

The mean amplitude and peak latency values of P1, N1, and P2 were analyzed. Following previous literature and visual inspection of the grand-averaged data, specific time windows and electrode clusters over the left and right occipital/parieto-occipital sites (X. Li et al., 2023) were identified for each component (P1, N1, and P2) separately. For each participant, the maximum or minimum response amplitude, depending on the polarity of the component, was first identified within the respective electrode cluster. For P1, the mean amplitude was extracted within the time window of 70–130 ms after stimulus onset. The time window of 190-250 was identified for P2. We determined N1 within the time window of 130–190 ms.

### 2.5 Multivariate pattern analysis

To investigate multivariate experimental differences between conditions within participant, the support vector machine (SVM) was employed to classify the segmented EEG signals under different stimulus conditions over the 125 scalp electrodes at each time point. Classification was performed in two ways: (1) a four-way classification to differentiate among all four conditions simultaneously, and (2) pairwise comparisons between conditions (2C-H vs. 2C-DH, 3C-H vs. 3C-DH, 2C-H vs. 3C-H, and 2C-DH vs. 3C-DH). All SVM analyses were conducted using MATLAB 2024b. The four-class classification training was conducted using the fitcecoc() function with error-correcting output codes, while binary classification training was implemented with the fitcsvm() function. For both approaches, model testing and prediction were executed using the predict() function.

Each epoch for decoding analysis was defined as −200 to 800 ms at the visual search display onset. Then the data was resampled to 100 Hz to increase the efficiency of the analysis. Variations in trial numbers across conditions were addressed by balancing the trial count for each condition within participants through subsampling of the available trials (Zhang, Carrasco, et al., 2024). For each participant, the trials in each condition were divided into 6 subsets (crossfolds), each containing an equal number of trials, which varied between 12 and 16 across conditions and participants due to differences in available trials. The single trials for each subset were then averaged in order to improve the signal-to-noise ratio (Bae & Luck, 2019; Zhang, Carrasco, et al., 2024), resulting in 6 averaged ERPs for each condition.

Decoding was performed using a leave-one-out cross validation approach with 6 crossfolds, where in each round, 5 averaged ERPs was selected for training classifier, and the remaining 1 averaged ERP from the same participant served as the testing set. This procedure was repeated 6tiems to ensure that each of the 6 averaged ERP was used as the test set once. To improve the precision of the decoding accuracy, the entire process was repeated 100 times, with random subsets of trials assigned to the averaged ERPs in each iteration (Zhang, Carrasco, et al., 2024; Zhang & Luck, 2025). The final decoding accuracy for each participant was calculated as the mean accuracy across 100 iterations.

### 2.6 Statistical analysis

Behavioral data and ERP components were analyzed using two-way repeated-measures analysis of variance (rmANOVA) to investigate the effects of color scheme (two-color vs. three-color) and color harmony (harmony vs. disharmony). The t-tests were further conducted to perform simple effects analyses with Bonferroni correction when significant interaction effects were identified in the behavioral or ERP results. Effect sizes were reported using partial eta squared (η_p_^2^) for ANOVA results and Cohen’s d for t-tests. The significance level was set at p<0.05.

For MVPA, the SVM was applied separately at each time point of the ERP waveforms, yielding time-resolved decoding accuracy for each participant. To evaluate whether the decoding accuracy significantly exceeded the chance level (0.25 for four-way classification, 0.5 for binary classification), one-tailed t-tests were performed at each time point. P-values were corrected for multiple comparisons using the false discovery rate (FDR) method.

## 3 Results

### 3.1 Behavior results

We conducted 2 (scheme complexity: Two-color vs. Three-color) × 2 (color harmony: Harmony vs. Disharmony) rmANOVA on the behavioral data. The results revealed a significant main effect of scheme complexity (F(1, 29) = 21.593, p <0.001, η_p_^2^ = 0.427), with scores in the two-color conditions significantly surpassing that in the three-color conditions. In contrast, the main effect of color harmony did not reach significance (F(1, 29) = 1.752, p = 0.196, η_p_^2^ = 0.057), suggesting that the harmony of the different color combinations had a limited impact on participants’ aesthetic evaluations. Additionally, the interaction between scheme complexity and color harmony was not significant (F(1, 29) = 1.764, p = 0.194, η_p_^2^ = 0.057). These findings emphasize the dominant role of scheme complexity in shaping participants’ aesthetic evaluations, with color harmony playing a comparatively limited role.

### 3.2 Univariate ERP results

#### 3.2.1 P1 component

As shown in Fig. 4, the P1 component exhibited a typical positive peak with a latency of approximately 99 ms across all four experimental conditions. The P1 amplitude for harmonious conditions was significantly larger than for disharmonious conditions, consistently observed across both two-color and three-color automotive schemes. The statistical analysis results revealed a significant main effect of the color harmony (F_(1,29)_ = 15.214, p = 0.001, η_p_^2^= 0.344) and a significant interaction effect between the two factors (F_(1,29)_ = 5.11, p = 0.031, η_p_^2^ = 0.15). However, we did not observe significant difference between the scheme complexity (F_(1,29)_ = 1.021, p = 0.321, η_p_^2^ = 0.034).

**Fig. 4.**
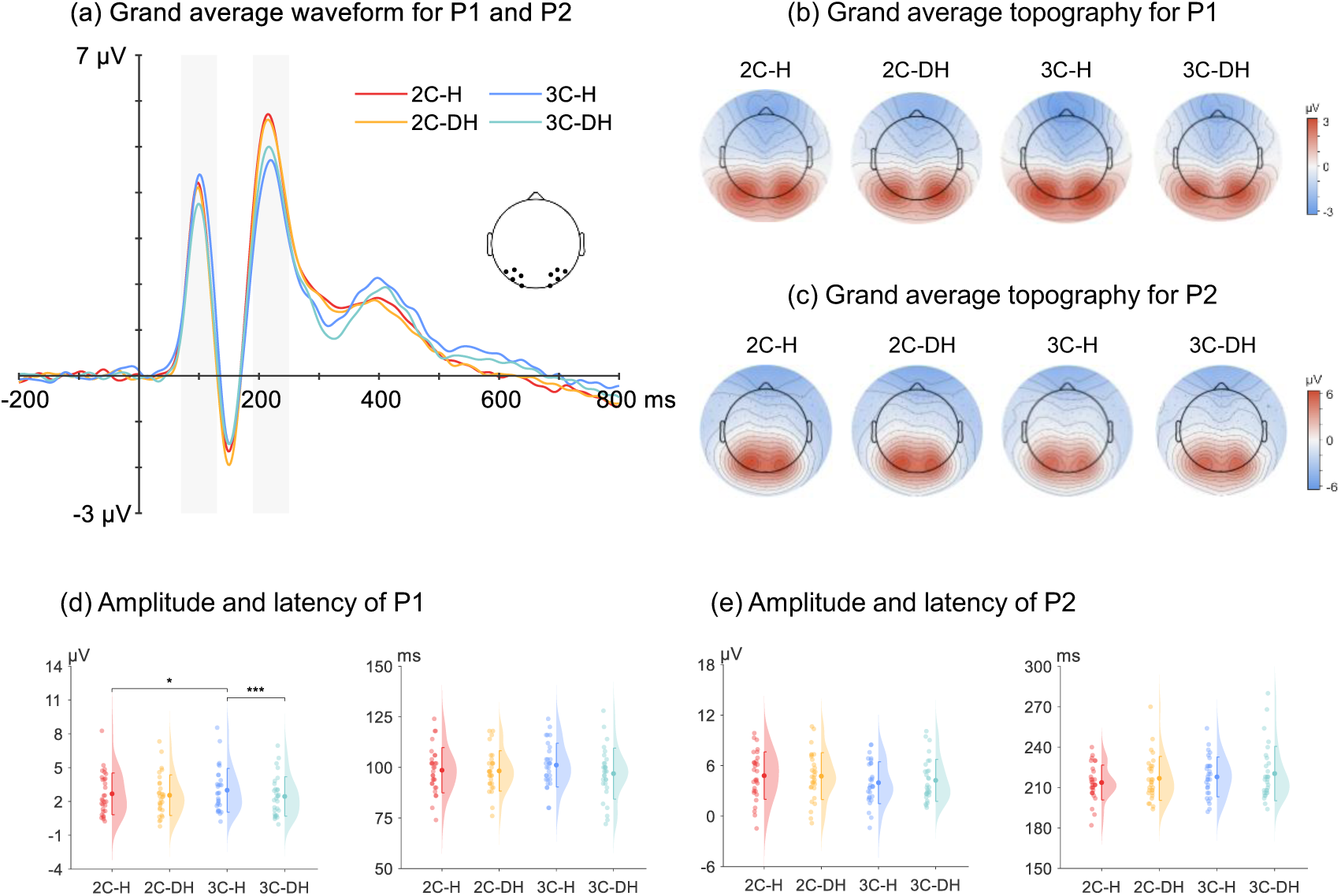
Grand average ERP waveforms (a), topographic maps (b), and raincloud plots (c) for P1 and P2 responses across all four conditions over a parieto-occipital cluster. The gray shading in the was the P1 time window (70-130 ms) and P2 time window (190-250 ms) used for calculating mean amplitude and latency values in the analysis. Panels (d) and (e) show the mean amplitude and latency values, with colored dots in the raincloud plots representing participant-level data and shaded areas illustrating the overall distribution, overlaid with the mean values.

Further simple effects analysis indicated a significant difference between harmonious and disharmonious conditions in the three-color combinations (t_(29)_ = 4.08, p < 0.001, Cohen’s d = 0.298) and between two-color and three-color combinations under harmonious conditions (t_(29)_ = −2.259, p = 0.032, Cohen’s d = −0.167). No significant effects were observed between different color combinations and harmony levels in other comparisons.

For P1 latency, the statistical analysis results showed no significant main effects of the Color factor or the Design factor, or interaction effect between the two factors.

#### 3.2.2 P2 component

The P2 component exhibited a positive peak at approximately 210 ms, with larger amplitudes in the two-color condition compared to the three-color condition. The statistical analysis results of P2 mean amplitude revealed a significant main effect of the scheme complexity factor on amplitude (F_(1,29)_ = 13.154, p = 0.001, η_p_^2^ = 0.312), whereas no significant effects were found either for the color harmony factor (F_(1,29)_ = 0.482, p = 0.493, η_p_^2^ = 0.016) or the interaction between scheme complexity and color harmony factors (F_(1,29)_ = 1.137, p = 0.295, η_p_^2^= 0.038).

For latency, the scheme complexity factor also showed a significant effect (F_(1,29)_ = 5.058, p = 0.032, η_p_^2^ = 0.149), whereas either the color harmony factor (F_(1,29)_ = 3.033, p = 0.092, η_p_^2^ = 0.095) or the interaction between scheme complexity and color harmony factors (F_(1,29)_ = 0.050, p = 0.824, η_p_^2^ = 0.002) was significant.

#### 3.2.3 N1 component

As illustrated in Fig. 5, the N1 component exhibited distinct differences across the four experimental conditions. Observations revealed that the N1 amplitude was larger in the 2C-H condition compared to the 2C-DH condition, while the latency was shorter in the 3C-DH condition compared to the 3C-H condition.

**Fig. 5.**
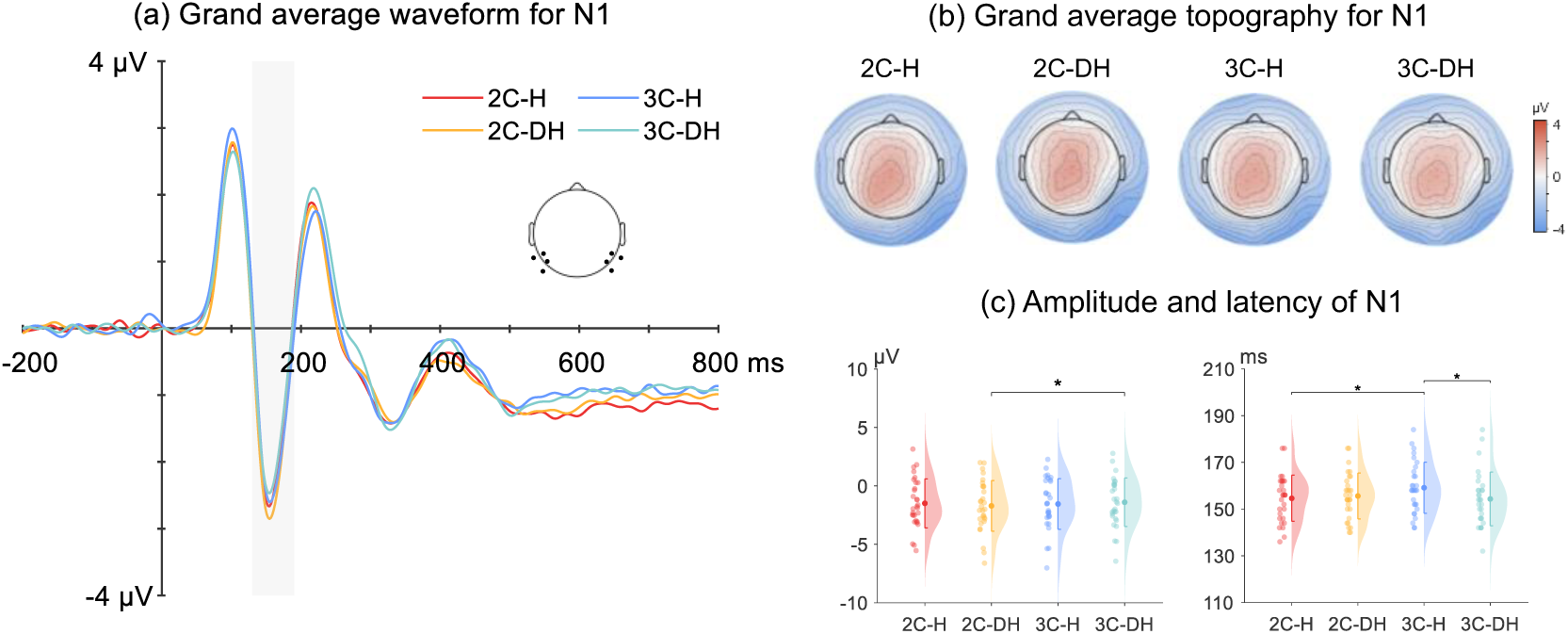
Grand average ERP waveforms (a), topographic maps (b), and raincloud plots (c) of N1 responses across all four conditions over a parieto-occipital cluster. The gray shading in the waveforms indicates the N1 time window (130-190 ms) used for calculating mean amplitude and latency values in the analysis. Raincloud plots (c) display mean amplitude and latency values, with colored dots representing participant-level data and shaded areas reflecting the overall distribution, overlaid with the mean values.

The N1 component results of amplitude showed no significant main effects for the scheme complexity (F_(1,29)_ = 2.022, p = 0.166, η_p_^2^ = 0.065) or the color harmony factor (F_(1,29)_ = 0.167, p = 0.686, η_p_^2^ = 0.006). However, a significant interaction between scheme complexity and color harmony was observed (F_(1,29)_ = 4.261, p = 0.048, η_p_^2^ = 0.128). Further simple effects analysis indicated a significant difference between two-color and three-color conditions under disharmonious conditions (t_(29)_ = −2.567, p = 0.016, Cohen’s d = 0.146).

For latency, there were no significant main effects of the scheme complexity factor (F_(1,29)_ = 3.644, p = 0.066, η_p_^2^ = 0.112) or the color harmony factor (F_(1,29)_ = 3.808, p = 0.061, η_p_^2^ = 0.116). However, a significant interaction effect between scheme complexity and color harmony was identified (F_(1,29)_ = 5.974, p = 0.021, η_p_^2^ = 0.171). Simple effects analysis revealed significant differences between harmony and disharmony in the three-color condition (t_(29)_ = 2.573, p = 0.015, Cohen’s d = 0.429) and between two-color and three-color conditions under harmony (t_(29)_ = −2.599, p = 0.015, Cohen’s d = −0.436), while no significant effects were observed in other comparisons.

### 3.3 Multivariate pattern analysis

To further investigate the differences among the conditions, we conducted MVPA for both all-condition classification and pairwise comparisons, including 2C-H vs. 2C-DH, 3C-H vs. 3C-DH, 2C-H vs. 3C-H, and 2C-DH vs. 3C-DH as shown in Fig. 6.

**Fig. 6.**
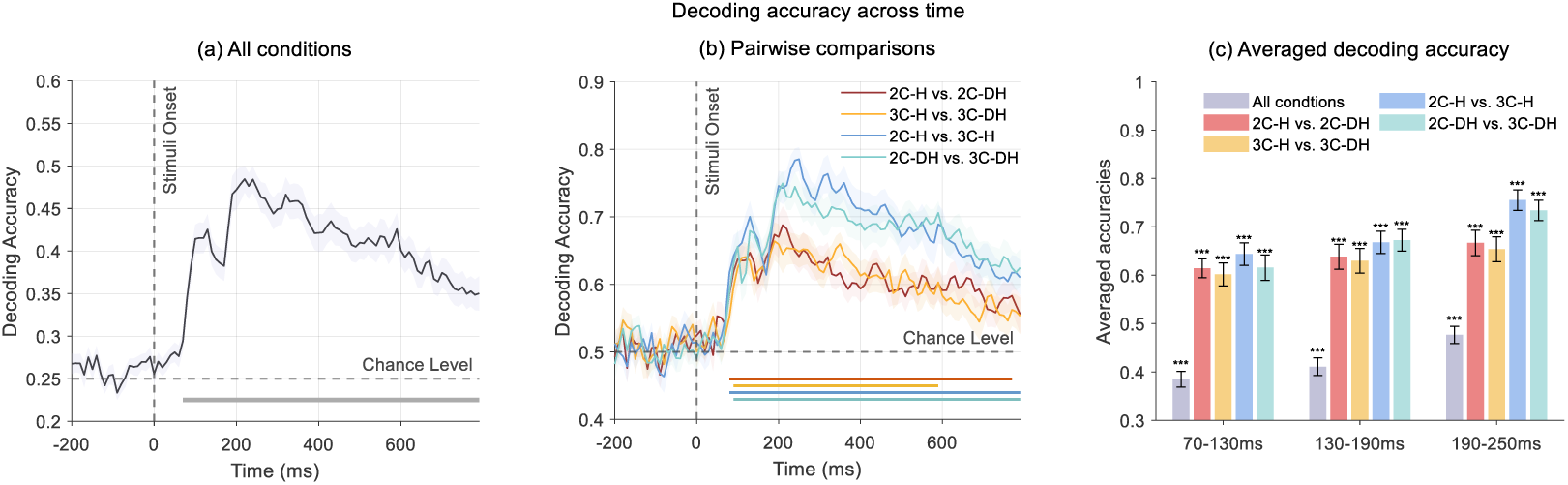
Decoding accuracy across time for all conditions and pairwise comparisons. (a) Decoding accuracy for all conditions. The solid line represents the mean decoding accuracy, and the shading indicates the ±1 standard error of the mean (SEM). The dashed horizontal line represents the chance level (0.25 for all conditions). (b) Decoding accuracy for pairwise comparisons: 2C-H vs. 2C-DH, 3C-H vs. 3C-DH, 2C-H vs. 3C-H, and 2C-DH vs. 3C-DH. Solid lines show the mean decoding accuracy for each comparison, and shaded areas represent the ±1 SEM. The dashed horizontal line represents the chance level (0.5 for pairwise comparisons). The corresponding horizontal and colored lines at the bottom indicate the time windows where decoding accuracies are significantly above chance level (FDR corrected p < 0.001) for the respective comparisons. (c) Averaged decoding accuracies within three time windows (70-130 ms, 130-190 ms, 190-250 ms). Bars represent mean accuracy per condition comparison with standard error; significance is indicated by asterisks (***p < .001). Accuracy is consistently above chance level across conditions and time windows.

The results demonstrated that the decoding accuracies for both all-condition and pairwise analyses were significantly above the chance level across nearly the entire time window after stimulus onset. Specifically, the significant time window for all-condition classification was 70-790 ms (see Fig. 6 a), whereas the significant time windows for the pairwise comparisons were 80-770 ms for 2C-H vs. 2C-DH, 90-590 ms for 3C-H vs. 3C-DH, 80-790 ms for 2C-H vs. 3C-H, and 90-790 ms for 2C-DH vs. 3C-DH, respectively (see Fig.6 b).

Notably, the behavioral data and P2 results indicate superior performance in two-color conditions compared to three-color conditions, a pattern that is also reflected in the decoding results. Specifically, decoding accuracy for two-color versus three-color comparisons was generally higher, aligning with the behavioral and P2 findings. While decoding accuracy for color harmony comparisons (2C-H vs. 2C-DH and 3C-H vs. 3C-DH) are relatively lower than that for scheme complexity comparisons (2C-H vs. 3C-H and 2C-DH vs. 3C-DH), it remained significantly above the chance level, particularly following stimulus onset and within the time windows of interest for ERP components. Decoding accuracies averaged within each time window revealed significantly above-chance performance across all comparisons.

Moreover, although scheme complexity showed no significant main effects on P1/N1, and color harmony had no significant effects on N1/P2, decoding analysis successfully classified both scheme complexity during P1/N1 windows and color harmony during N1/P2 windows. These findings underscore the sensitivity of MVPA in revealing subtle neural differences that may not be captured by traditional ERP analyses.

## 4 Discussion

We explored whether the aesthetic values calculated based on color harmony correspond to users’ neurophysiological responses, and whether such correspondence occurs across perceptual and evaluative dimensions. Specifically, we analyzed the effects of color harmony and scheme complexity on subjective aesthetic ratings and neural activity with ERPs and MVPA. Our findings indicated that scheme complexity significantly influenced behavioral ratings, whereas color harmony had a more pronounced impact on neural responses. Moreover, MVPA decoding substantiated the dissociable neural representations between color harmony and scheme complexity conditions, aligning with the stage-specific ERP components observed. Consistent with our hypothesis, the results suggest that scheme complexity plays a more dominant role in late-stage neural processing (reflected in the P2 component) and behavioral outcomes, whereas color harmony elicits distinct neural signatures during early stages, particularly in the P1 and N1 components. This highlights the importance of distinguishing cognitive stages in aesthetic perception models.

The behavioral data revealed that the scheme complexity significantly influenced users’ aesthetic ratings, with two-color schemes receiving higher scores than for three-color schemes. This means that less complex information is preferable and easier to process according to the fluency theory of aesthetics (Reber et al., 2004). However, color harmony did not have a significant effect on subjective ratings or interest scores. This result is consistent with previous findings that color harmony does not fully determine individuals’ preferences for color combinations (Ou et al., 2004b). Additionally, previous research has suggested that color preference is shaped by individual experience, cultural background, and other contextual factors (Palmer & Schloss, 2010). This complexity may explain why color harmony did not significantly impact subjective ratings or interest scores in the present study, indicating that aesthetic evaluation is likely influenced by multiple interacting factors beyond harmony alone.

The ERP results indicated that color harmony had a significant effect on P1 amplitude, with harmonious color combinations eliciting greater P1 amplitudes than for disharmonious color combinations. The P1 amplitude enhancement for harmonious stimuli was particularly prominent in three-color combinations, indicating that both color harmony and chromatic complexity modulate early visual processing. This finding aligns with predictions from color harmony model and with previous ERP studies on symmetry aesthetics (Höfel & Jacobsen, 2007; Makin et al., 2012; Moon & Spencer, 1944a), which show that the visual system rapidly responds to aesthetic information at low-level processing stages. Moreover, P1 is highly sensitive to the intensity of attention allocated to stimuli, with its amplitude increasing as attention strengthens (Smith et al., 2003), and it is typically associated with physical stimulus features (Perri et al., 2019). This further suggests that in more complex three-color combinations, harmonious color schemes may enhance visual attention, thereby influencing early perceptual processing. However, P1 latency did not show significant effects, indicating that color harmony primarily modulates the strength of neural responses rather than processing speed.

The amplitude and latency of P2 component were primarily influenced by scheme complexity, with two-color schemes eliciting greater P2 amplitude and shorter latencies compared to three-color schemes. This finding aligns with behavioral data, which showed that users rated two-color schemes higher than three-color schemes. According to Gestalt psychology and Berlyne’s aesthetic theory, visual complexity theory predicts that the complexity of stimuli influences arousal potential and hedonic evaluation (valence) in an inverted U-shaped pattern (Berlyne, 1974; Lazard & King, 2020). Within this framework, the increased complexity of three-color schemes raises visual cognitive load, leading to reduced P2 resource allocation, thereby decreasing P2 amplitude and prolonging latency. Similarly, Liu et al., 2022 also found that low-complexity webpages elicited greater P2 amplitudes than high-complexity webpages, further suggesting that lower visual complexity facilitates cognitive resource allocation and emotional arousal. However, color harmony did not significantly affect P2, and no interaction effect between scheme complexity and color harmony was observed.

For N1 component, we observed that harmonious two-color schemes elicited greater N1 amplitude than disharmonious ones. This aligns with visual aesthetic preference research, which suggests that preferred stimuli tend to elicit greater N100 amplitude (Guo et al., 2022; W. Liu et al., 2022). This also supports the view that harmonious color combinations receive greater perceptual resource allocation during early stages of visual processing as reported in the previous study (Kok et al., 2012). On the other hand, the latency of N1 for disharmonious three-color schemes was shorter than for harmonious three-color schemes, suggesting that violations of color harmony may trigger rapid prediction error signals within the visual system (Friston, 2005). Such early error detection may initiate subsequent conflict-monitoring processes (Botvinick et al., 2001), reflecting the brain’s sensitivity to irregularities in complex color input at an early perceptual level.

The MVPA results not only aligned closely with behavioral data but also revealed neural distinctions that traditional ERP statistical analyses did not detect. While behavioral results showed a significant difference between two-color and three-color conditions but no significant effect of harmony or interaction, MVPA decoding accuracy was significantly above chance level from approximately 70 ms to 600 ms post-stimulus for all pairwise comparisons. In particular, the comparisons between two-color and three-color conditions reached higher decoding accuracy than the comparisons between harmony and disharmony conditions, which is consistent with the behavioral results and the P2 findings. Additionally, although ERP results showed no significant differences between two-color and three-color conditions in the P1 and N1 components, MVPA decoding accuracy was still significantly above chance level in these time windows, suggesting that neural signals carried distinguishable information. Similarly, while N1 and P2 amplitudes did not show significant differences between harmony and disharmony conditions, MVPA decoding accuracy remained significantly above chance level, indicating that harmony-related neural differences were still present. These results suggest that MVPA can detect neural distinctions that are not evident in traditional ERP amplitude analyses, providing a complementary approach for studying visual processing.

There are still some limitations of this study that should be further considered. First, the lack of assessment for red-green color blindness may have influenced the results. Although participants self-reported normal color vision, subtle differences could still affect neural responses to color harmony. Future studies should use standardized tests to ensure consistency. Second, the focus on automotive color schemes may limit generalizability. Expanding research to UI design, interior decor, and fashion could provide broader insights. Additionally, individual differences in color perception and aesthetic preference, including personal color preferences and cultural background, were not controlled. Future studies should incorporate color preference assessments and individual-level neural analyses to explore variations in color harmony processing across populations.

## 5 Conclusion

This study employed traditional univariate analysis and MVPA to examine whether the aesthetic values calculated based on color harmony theory reflect users’ neurophysiological response patterns. ERP results showed that P1 and N1 were primarily influenced by color harmony, while P2 was mainly affected by scheme complexity. MVPA results further demonstrated that the neural system effectively distinguishes scheme complexity and color harmony, with scheme complexity being successfully decoded in the P1 and N1 time windows, while color harmony exhibited classification effects in the N1 and P2 windows. Compared to traditional ERP analysis, MVPA provided greater sensitivity in revealing the easier processing of two-color schemes over three-color schemes and confirmed the influence of color harmony on early visual processing.

## List of abbreviations

ERP: Event-related potential
MVPA: Multivariate pattern analysis
rmANOVA: Repeated-measures analysis of variance
SVM: Support vector machine
2C-DH: Two-color disharmony
2C-H: Two-color harmony
3C-DH: Three-color disharmony
3C-H: Three-color harmony.

## Declarations

### Ethics approval and consent to participate

The research protocol was carried out in accordance with the Declaration of Helsinki and was approved by the Ethics Committee of the University of Jyväskylä (approval ID: l74/13.00.04.00/2024). Informed consent was obtained from all participants prior to their involvement in the study, ensuring their understanding and willingness to participate.

### Consent for publication

The findings presented in this manuscript are original and have neither been published elsewhere nor submitted for consideration to any other publisher by any of the authors.

### Data availability

The datasets generated and/or analyzed during this study are available from the corresponding author upon reasonable request.

## Acknowledgements

Not applicable.

## Contributions

L.H. conceptualized the study, curated and analyzed the data, developed the methodology, created visualizations, and wrote the original draft. G.Z., T.P., F.C., and T.K. contributed to visualization and manuscript review and editing. X.L., D.T., T.P., and F.C. contributed to conceptualization. T.K. also led project administration and supervision. All authors reviewed and approved the final manuscript.

## References

Bae, G.-Y., & Luck, S. J. (2019). Decoding motion direction using the topography of sustained ERPs and alpha oscillations. Neuroimage, 184, 242–255.

Berlyne, D. E. (1974). Studies in the new experimental aesthetics: Steps toward an objective psychology of aesthetic appreciation. Hemisphere.

Botvinick, M. M., Braver, T. S., Barch, D. M., Carter, C. S., & Cohen, J. D. (2001). Conflict monitoring and cognitive control. Psychological Review, 108(3), 624.

Burchett, K. E. (2002). Color harmony. Color Research & Application, 27, 28–31.

Cheng, Y. (2018). Study of electroencephalography cognitive model of product image. Journal of Mechanical Engineering, 54(23), 126.

Chuang, M. C., & Ou, L. C. (2001). Influence of a holistic color interval on color harmony. Color Research & Application, 26(1), 29–39.

Delorme, A., & Makeig, S. (2004). EEGLAB: an open source toolbox for analysis of single-trial EEG dynamics including independent component analysis. Journal of Neuroscience Methods, 134(1), 9–21.

Ding, M., Ding, T., Chen, X., & Shi, F. (2022). Using event-related potentials to identify user’s moods induced by product color stimuli with different attributes. Displays, 74.

Ding, M., Qin, K., Qin, H., & Sun, M. (2023). Using event-related potentials to identify user emotion caused by product color attribute. Displays, 79, 102460.

Ding, M., Song, M., Pei, H., & Cheng, Y. (2021). The emotional design of product color: An eye movement and event-related potentials study. Color Research & Application, 46(4), 871–889.

Eimer, M. (2014). The time course of spatial attention: insights from event-related brain potentials. The Oxford Handbook of Attention, 1, 289–317.

Friston, K. (2005). A theory of cortical responses. Philosophical Transactions of the Royal Society B: Biological Sciences, 360(1456), 815–836.

Guo, F., Li, M., Chen, J., & Duffy, V. G. (2022). Evaluating users’ preference for the appearance of humanoid robots via event-related potentials and spectral perturbations. Behaviour & Information Technology, 41, 1381–1397.

Herbert, C., Kissler, J., Junghofer, M., Peyk, P., & Rockstroh, B. (2006). Processing of emotional adjectives: Evidence from startle EMG and ERPs. Psychophysiology, 43, 197–206.

Hillyard, S. A., & Anllo-Vento, L. (1998). Event-related brain potentials in the study of visual selective attention. Proceedings of the National Academy of Sciences, 95(3), 781–787.

Höfel, L., & Jacobsen, T. (2007). Electrophysiological indices of processing aesthetics: Spontaneous or intentional processes? International Journal of Psychophysiology, 65(1), 20–31.

Hsiao, S.-W., Chiu, F.-Y., & Chen, C. S. (2008). Applying aesthetics measurement to product design. International Journal of Industrial Ergonomics, 38(11–12), 910–920.

Hsiao, S.-W., & Yang, M.-H. (2017). A methodology for predicting the color trend to get a three-colored combination. Color Research & Application, 42(1), 102–114.

Jia, Y., & Tyler, C. W. (2019). Measurement of saccadic eye movements by electrooculography for simultaneous EEG recording. Behavior Research Methods, 51, 2139–2151.

Kok, P., Jehee, J. F., & De Lange, F. P. (2012). Less is more: expectation sharpens representations in the primary visual cortex. Neuron, 75(2), 265–270.

Lazard, A. J., & King, A. J. (2020). Objective design to subjective evaluations: Connecting visual complexity to aesthetic and usability assessments of eHealth. International Journal of Human-Computer Interaction, 36(1), 95–104.

Li, K.-R., Zheng, Z.-Q., Wang, P.-H., & Yan, W.-J. (2022). Research on the colour preference and harmony of the two-colour combination buildings. Color Research & Application, 47, 980–991.

Li, X., Vuoriainen, E., Xu, Q., & Astikainen, P. (2023). The effect of sad mood on early sensory event-related potentials to task-irrelevant faces. Biological Psychology, 178, 108531.

Liu, B., Meng, X., Wu, G., & Huang, Y. (2012). Feature precedence in processing multifeature visual information in the human brain: an event-related potential study. Neuroscience, 210, 145–151.

Liu, W., Cao, Y., & Proctor, R. W. (2022). The roles of visual complexity and order in first impressions of webpages: an ERP study of webpage rapid evaluation. International Journal of Human-Computer Interaction, 38(14), 1345–1358.

Luck, S. J., Heinze, H., Mangun, G., & Hillyard, S. A. (1990). Visual event-related potentials index focused attention within bilateral stimulus arrays. II. Functional dissociation of P1 and N1 components. Electroencephalography and Clinical Neurophysiology, 75(6), 528–542.

Luck, S. J., & Hillyard, S. A. (1994). Electrophysiological correlates of feature analysis during visual search. Psychophysiology, 31, 291–308.

Makin, A. D., Wilton, M. M., Pecchinenda, A., & Bertamini, M. (2012). Symmetry perception and affective responses: A combined EEG/EMG study. Neuropsychologia, 50(14), 3250–3261.

Moon, P., & Spencer, D. E. (1944a). Aesthetic measure applied to color harmony. Journal of the Optical Society of America, 34(4), 234–242.

Moon, P., & Spencer, D. E. (1944b). Geometric formulation of classical color harmony. Journal of the Optical Society of America, 34(1), 46–59.

Omoto, S., Kuroiwa, Y., Otsuka, S., Baba, Y., Wang, C., Li, M., Mizuki, N., Ueda, N., Koyano, S., & Suzuki, Y. (2010). P1 and P2 components of human visual evoked potentials are modulated by depth perception of 3-dimensional images. Clinical Neurophysiology, 121, 386–391.

Ou, L.-C., & Luo, M. R. (2006). A colour harmony model for two-colour combinations. Color Research & Application, 31(3), 191–204.

Ou, L.-C., Luo, M. R., Woodcock, A., & Wright, A. (2004a). A study of colour emotion and colour preference. Part I: Colour emotions for single colours. Color Research & Application, 29, 232–240.

Ou, L.-C., Luo, M. R., Woodcock, A., & Wright, A. (2004b). A study of colour emotion and colour preference. part II: colour emotions for two-colour combinations. Color Research & Application, 29(4), 292–298.

Palmer, S. E., & Schloss, K. B. (2010). An ecological valence theory of human color preference. Proceedings of the National Academy of Sciences, 107(19), 8877–8882.

Perri, R. L., Berchicci, M., Bianco, V., Quinzi, F., Spinelli, D., & Di Russo, F. (2019). Perceptual load in decision making: The role of anterior insula and visual areas. An ERP study. Neuropsychologia, 129, 65–71.

Reber, R., Schwarz, N., & Winkielman, P. (2004). Processing fluency and aesthetic pleasure: Is beauty in the perceiver’s processing experience? Personality and Social Psychology Review, 8(4), 364–382.

Schloss, K. B., & Palmer, S. E. (2011). Aesthetic response to color combinations: preference, harmony, and similarity. *Attention, Perception*, & Psychophysics, 73, 551–571.

Scott, G. G., O’Donnell, P. J., Leuthold, H., & Sereno, S. C. (2009). Early emotion word processing: Evidence from event-related potentials. Biological Psychology, 80(1), 95–104.

Shih-Wen, H., Yang, M.-H., & Lee, C.-H. (2017). An aesthetic measurement method for matching colours in product design. Color Research \& Application, 42(5), 664–683.

Smith, N. K., Cacioppo, J. T., Larsen, J. T., & Chartrand, T. L. (2003). May I have your attention, please: Electrocortical responses to positive and negative stimuli. Neuropsychologia, 41(2), 171–183.

Zhang, G., Carrasco, C. D., Winsler, K., Bahle, B., Cong, F., & Luck, S. J. (2024). Assessing the effectiveness of spatial PCA on SVM-based decoding of EEG data. NeuroImage, 293, 120625.

Zhang, G., Garrett, D. R., & Luck, S. J. (2024a). Optimal filters for ERP research I: A general approach for selecting filter settings. Psychophysiology, 61(6), e14531.

Zhang, G., Garrett, D. R., & Luck, S. J. (2024b). Optimal filters for ERP research II: Recommended settings for seven common ERP components. Psychophysiology, 61(6), e14530.

Zhang, G., Garrett, D. R., Simmons, A. M., Kiat, J. E., & Luck, S. J. (2024). Evaluating the effectiveness of artifact correction and rejection in event-related potential research. Psychophysiology, 61(5), e14511.

Zhang, G., & Luck, S. (2025). Assessing the impact of artifact correction and artifact rejection on the performance of SVM-based decoding of EEG signals. NeuroImage, 121304.

