## Supplementary material for "EEG-based analyses reveal different temporal processing patterns in aesthetic evaluation"

Limin Hou^1,2,3,4^, Guanghui Zhang^1,2,5^*, Xiawen Li^6^, Dong Tang^7^, Tiina Parviainen^8^,

Fengyu Cong^3,4,9^, Tommi Kärkkäinen^4^

*^1^* *Institute of Psychological and Brain Sciences, Liaoning Normal University, Dalian, Liaoning, 116029, China*

*^2^* *Key Laboratory of Brain and Cognitive Neuroscience, Liaoning Province, Dalian, 116029, China*

*^3^* *School of Biomedical Engineering, Faculty of Medicine, Dalian University of Technology, Dalian, 116024, Liaoning Province, China*

*^4^ Faculty of Information Technology, University of Jyväskylä, P.O.Box FI-40014 Jyväskylä, Finland*

*^5^* *Center for Mind and Brain, University of California–Davis, Davis, CA, 95618, USA*

*^6^ Department of Physical Education, Shanghai University of Medicine and Health Sciences, Shanghai, 201318, China*

*^7^* *School of Foreign Studies, China University of Petroleum (East China), Qingdao, 266580, China*

*^8^ Centre for Interdisciplinary Brain Research, Department of Psychology, University of Jyväskylä, P.O.Box FI-40014 Jyväskylä, Finland*

*^9^ Key Laboratory of Social Computing and Cognitive Intelligence (Dalian University of Technology), Ministry of Education, Dalian, Liaoning 116024, China*

*Corresponding author: Guanghui Zhang

Research Center of Brain and Cognitive Neuroscience, Liaoning Normal University, Dalian, Liaoning, 116029, China..

In the supplementary materials, we provide the selection process for identifying the most user-preferred schematic layouts of two-tone and three-tone car designs.

Based on the typical classification of body panels in modern automobiles (Doshi et al., 2024; Gauhar et al., 2017), the car body in this study was divided into six major regions: Zone 1 (front hood), Zone 2 (bumper), Zone 3 (front and rear fenders), Zone 4 (doors), Zone 5 (trunk lid), and Zone 6 (roof) (see Fig. S1a). Two-tone and three-tone layout schemes were generated by combining any two or three of these zones, respectively. Informed by extensive observation of existing and conceptual multicolor vehicle schemes, seven distinct layouts were developed for the two-tone condition (see Fig. S1b), and nine for the three-tone condition (see Fig. S1c). To eliminate the influence of specific color preferences, all layout schemes were initially rendered using neutral tones.

A preference survey was then conducted to evaluate participants’ aesthetic preferences for each layout. The questionnaire was designed to assess participants’ evaluations of two-tone and three-tone car body schemes, as well as their general attitudes toward multicolor design. Administered online via Microsoft Forms, the survey consisted of four sections: (1) demographic and background information; (2) preference ratings for seven two-tone car design schemes; (3) preference ratings for nine three-tone design schemes; and (4) follow-up questions regarding participants’ perceptions, preferences, and expectations related to multicolor layouts. All items were presented in randomized order, and preference ratings were measured using a 5-point Likert scale ranging from 1 (strongly dislike) to 5 (strongly like). A total of 118 valid responses were collected through the online platform. Participants were recruited voluntarily, and informed consent was obtained prior to participation. Only fully completed questionnaires were included in the final analysis.

The sample included 57 males and 61 females, with 49% aged 18–25 and 47% aged 26-35. Additionally, 62% of participants reported prior driving experience. Regarding attitudes toward car color, 40% of participants considered color to be highly important in car design, and 48% believed that car color influences their purchasing decisions. Notably, 95% of respondents expressed a preference for customizable color options, 94% had encountered multicolor cars in real life, and 77% regarded multicolor car design as a reflection of innovation and individuality.

Based on participants’ preference ratings, the third layout in the two-tone condition and the sixth layout in the three-tone condition were identified as the most preferred by users. The average preference ratings for all two-tone and three-tone car body layout schemes are shown in Fig. S2.

**Figure S1**. Selection process and user-preferred schematic layouts of two-tone and three-tone car designs. (a) Six major body regions. (b) Seven different two-tone combination designs. (c) Nine different three-tone combination designs.

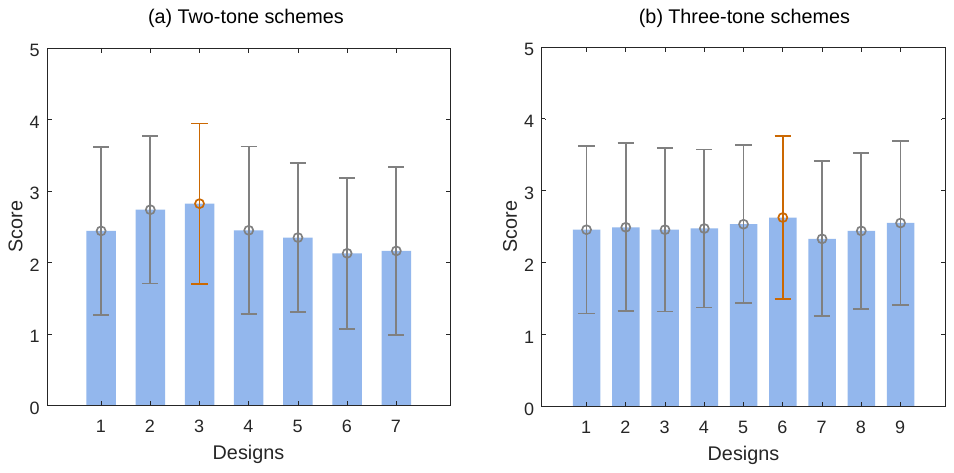

**Figure S2**. Mean preference ratings for car body color designs. (a) Two-tone designs: scheme 3 rated highest. (b) Three-tone designs: scheme 6 rated highest.

**Reference**

Doshi, S., Vaghosi, K., & Mehta, N. (2024). Progress in the welding of AL alloy thin sheet and future prospectus for automobile. *ITEGAM-JETIA*, *10*(46), 42–49.

Gauhar, N., Palm, C., & Vollmer, R. (2017). Line Concept For High Volume Production O f Hot-Formed High-Strength Aluminium Alloys. *European Aluminium Congress*.
